# Floral Radiometry: A Biophysical Basis to Characterize Landscapes

**DOI:** 10.1101/088583

**Authors:** R. Jaishanker, N.P. Sooraj, M. Somasekheran Pillai, K. Athira, Ammini Joseph, Aswathy Krishna, I Indu, V SarojKumar

## Abstract

This paper presents early results on studies of floral spectral reflectance of angiosperms and introduces floral radiometry as an emergent dimension of ecological research in India. Floral spectral reflectance of 121 angiosperm species was measured using hand-held spectroradiometer. The authors describe spectral reflectance of seven representative species within 350-800 nanometer region of the electromagnetic spectrum. Characteristic absorption and reflection of flowers, in ultraviolet and visible regions of the spectrum is reported. Near infrared reflectance was consistently high for all species studied. Flower color is unique to a species and importance of understanding flower color from pollinator’s perspective is highlighted.

## 1. Introduction

Flowers are *Bill boards* that plants use to attract pollinators. Floral coloration is the result of combined effect of light scattering by irregular cell complexes and wavelength selective absorbing pigments. Incoherently scattered light in the complementary wavelength determines the color of petals^1^. Deciphering flower color from a pollinator perspective is pivotal to floral biology, pollination studies and landscape ecology^2-5^. Floral radiometry presents a straightforward biophysical means to characterize landscapes and monitor temporal changes.

Potentials of radiometric studies of flowers were extensively discussed by Sir C.V. Raman during the 1960s^6-9^. Whilst extensive studies have been carried out on flower color and plant pigments using light transmission technique (spectroscopy)^10-13^ there hardly exist any study on floral reflection in India. However numerous studies have been reported internationally^14-18^. Here the authors present the findings of the first radiometric studies of flowers in India.

## 2. Study area and Methodology

The study was carried out in Trivandrum, Kerala, India. Floral spectral reflection of 121 species of flowering angiosperms was measured in a dedicated campaign during February-March 2016. Situated between 8°17’N and 8°54’N latitudes and 76°41’E and 77°17’ E longitudes, Trivandrum enjoys a tropical climate. The region falls in the Malabar phytogeographical province^19^ and harbors over 5000 species of flowering plants spread across 1500 genera and over 200 Families^20&21^.

Floral spectral reflectance were measured using hand-held spectroradiometer (Analytical Spectral Device, Inc. ASD Inc. Field spec^®^ 3)^22^. It records continuous reflected spectra from 350 nm to 2500 nm. All measurements were taken in fully open, healthy flowers. The field campaign was intentionally done on sunny days between 10:00 and 11:30 AM. Each floral spectral value recorded was a mean of 50 iterations aggregated across 15 samples of each species. Standard white reference correction was performed for each sample before and after recording. All reflectance spectra were processed using ASD View spec Pro for further analysis.

Quantitative analysis of spectral reflectance of representative violet, blue, yellow, orange, red, pink and white colored flowers were carried out using R stat. Details of the representative flowers selected for this study are indicated in Table 1. Human perceived colors of flowers used for the study are illustrated in Figure 1. Spectral behavior of the representative flowers in Ultraviolet (UV); 350-399 nm, Blue (440-489 nm), Green (490-569 nm), Yellow (570-585 nm), Red (620-699 nm) and Near infrared (NIR) (700-800 nm) regions of electromagnetic spectrum constitutes findings of this study.

**Figure 1 (a-g).**
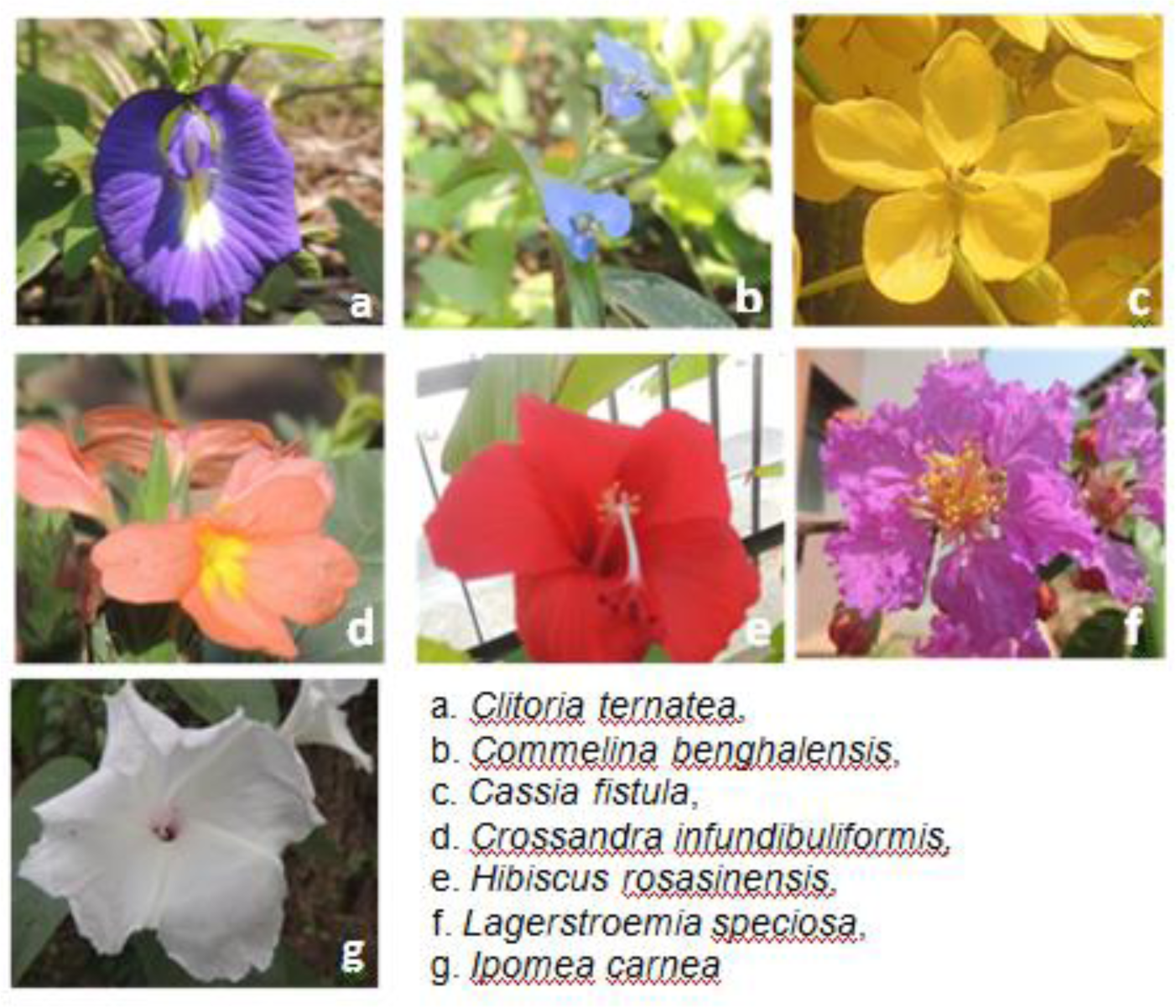
Species name and human perceived color of flowers

**Table 1.**
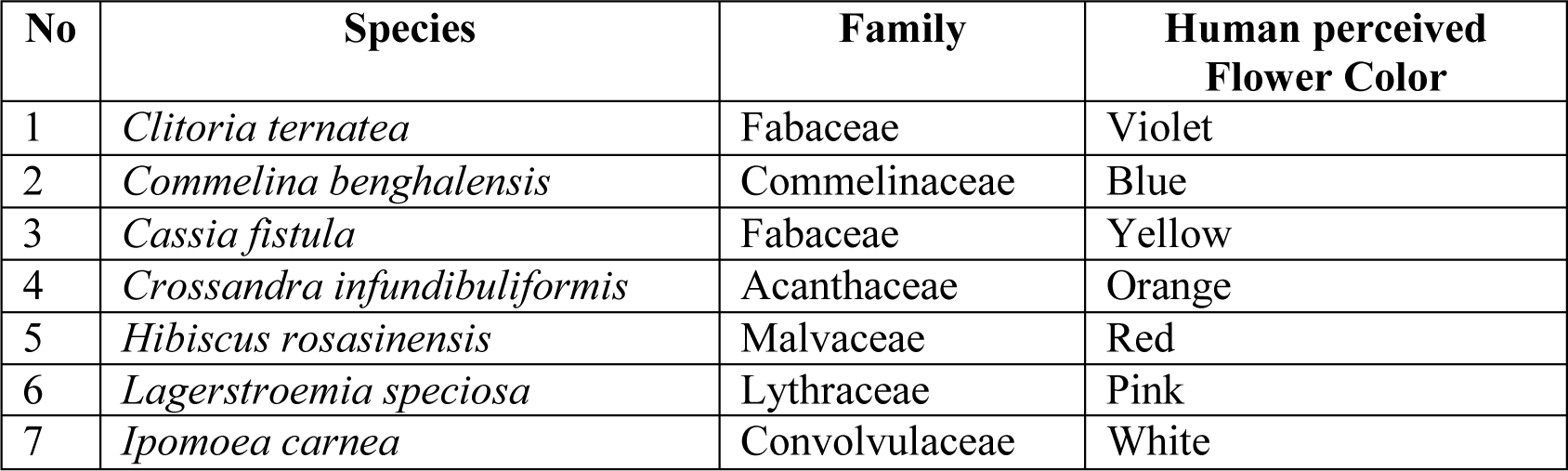
Details of selected flowers used for the present study

## 3. Results and Discussion

Table 2. summarizes the floral spectral reflectance of the seven species listed in Table 1. Characteristic floral spectral response curve of angiosperms used in the present study is depicted in Figure 2. It is interesting to note that all the seven flowers are highly reflective in the NIR region. At 28% *Clitoria ternatea* has the least and *Lagerstromia speciosa* (70%), the highest NIR reflectance. Whilst strong UV absorption (98%) was observed in *Clitoria ternatea*, *Cassia fistula* (20%) and *Cossandra infundibuliformis* (17%) reported high UV reflection. However UV reflection of *Cossandra infundibuliformis* showed a rising trend from 350 to 399 nm and *Cassia fistula* showed a decreasing trend.

**Figure 2.**
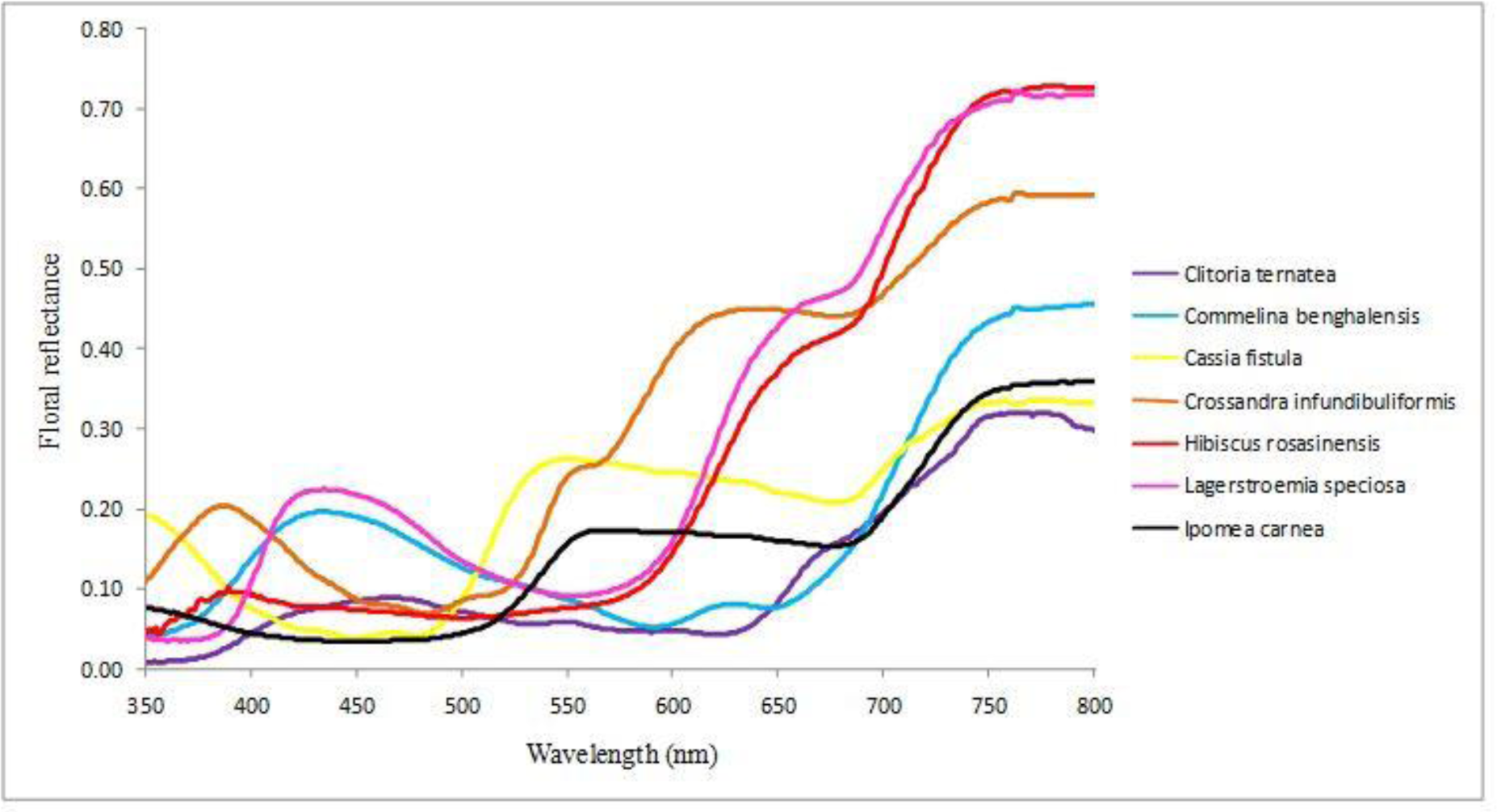
Spectral response curves of flowers

**Table 2.** Spectral region wise statistical parameters of floral reflectance

| Species | Spectral Region | Mean Floral Reflection (%) (Min. Max) | SD (%) | Skewness z score |
| --- | --- | --- | --- | --- |
| *Clitoria ternatea* | UV | 1.8 (0.8, 4.3) | 1 | 3.080* |
|  | Blue | 8.5 ( 7.8, 8.9) | 0.3 | -2.080* |
|  | Green | 6.1 (5, 7.7) | 0.7 | 2.468 |
|  | Yellow | 4.8 (4.8, 4.9) | 0.05 | 1.488 |
|  | Red | 11.3 (4.4, 19.5) | 5.2 | -0.108 |
|  | NIR | 28.2 (19.7, 32) | 3.7 | -3.625* |
| *Commelina benghalensis* | UV | 7.3 (4.2, 13.6) | 2.9 | 2.214* |
|  | Blue | 17.4 (14.2, 19.6) | 1.7 | -1.202 |
|  | Green | 10.4 (6.9, 14.1) | 1.9 | 0.175 |
|  | Yellow | 6.1 (5.5, 6.8) | 0.4 | 0.539 |
|  | Red | 11.2 (7.7, 21.8) | 4.1 | 4.123* |
|  | NIR | 39.8 (22.4, 45.6) | 6.9 | -4.733* |
| *Cassia fistula* | UV | 14 (7.7, 19.3) | 3.8 | -0.448 |
|  | Blue | 4.3 (3.9, 5.4) | 0.4 | 3.567* |
|  | Green | 19.8 (5.6, 26.27) | 7.2 | -2.799 |
|  | Yellow | 25.4 (25.1, 25.74) | 0.2 | 0.323 |
|  | Red | 22.2 (20.8, 24.76) | 1 | 0.937 |
|  | NIR | 31.7 (25, 33.55) | 2.4 | -5.246* |
| *Crossandra infundibuliformis* | UV | 17.3 (10.9, 20.5) | 3 | -2.234* |
|  | Blue | 8.1 (6.9, 10.3) | 0.9 | 2.499* |
|  | Green | 15.6 (7.3, 26.6) | 7.12 | 1.401 |
|  | Yellow | 29.8 (26.9, 33.2) | 2.1 | 0.246 |
|  | Red | 44.8 (44.1, 46.9) | 0.5 | 5.699* |
|  | NIR | 56.3 (47.1, 59.6) | 3.9 | -4.379* |
| *Hibiscus rosasinensis* | UV | 7.5 (4, 10.2) | 1.9 | -1.142 |
|  | Blue | 7.1 (6.5, 7.5) | 0.3 | -0.395 |
|  | Green | 7 (6.3, 8.3) | 0.6 | 1.316 |
|  | Yellow | 9.2 (8.4, 10.3) | 0.6 | 0.537 |
|  | Red | 38.1 (24.2, 49.5) | 6.2 | -1.855 |
|  | NIR | 67.7 (50.1, 72.8) | 6.7 | -5.096* |
| *Lagerstroemia speciosa* | UV | 4.7 (3.5, 10.2) | 1.7 | 5.558* |
|  | Blue | 19.8 (15.5, 22.4) | 2.1 | -1.650 |
|  | Green | 11 (9.2, 15.3) | 1.8 | 3.175 |
|  | Yellow | 10.7 (9.8, 11.7) | 0.6 | 0.645 |
|  | Red | 43.5 (28.3, 54.9) | 6.3 | -2.175 |
|  | NIR | 68.4 (55.6, 72.2) | 4.6 | -5.642* |
| *Ipomea carnea* | UV | 6.1 (4.5, 7.6) | 1 | -0.510 |
|  | Blue | 3.5 (3.4, 3.9) | 0.1 | 3.674* |
|  | Green | 10.1 (3.9, 17.2) | 5 | 0.758 |
|  | Yellow | 17.1 (17, 17.2) | 0.05 | -1.626 |
|  | Red | 16.1 (15.3, 18.9) | 0.8 | 5.413* |
|  | NIR | 31.6 (19.2, 35.9) | 5.2 | -4.338* |

\* indicates skewed distribution

* indicates skewed distribution

*Commelina benghalensis* is a true blue reflecting flower. *Lagerstroemia speciosa* flower that appears pink to human observer absorbs almost equally green, yellow (89% and 90%) and absorbs slightly less in blue (79%). It has a higher absorption in red (56.5%). Human perceived color of *Lagerstroemia speciosa* is the complementary analogue resulting from absorption of blue, green and yellow. *Hibiscus rosasinensis* is characterized by its exponential increase in red reflection. It shows a mean reflection of 40% in red region. Flowers of *Ipomoea carnea* exhibits low to modest reflection across the entire spectrum studied.

It is evident from Table 2 that 60% of the flowers studied show a positive skew in their reflectance at lower end of spectrum. NIR reflectance in all species was skewed. Green and yellow floral reflectance of all species studied was normally distributed. Incoherently scattered light in the complementary wavelength determines color of flowers^1^. Flowers with blue absorbing carotenoid appear yellow and those with blue absorbing anthocyanin appear purple^23^. Specific reflection and absorption by flowers in the UV region indicates to the presence of chalcones/ alurones class of flavonoids^24^. Importance of UV absorption and reflection pattern in flowers was reported as early as 189415. Specific pattern of floral reflectance (iridescence) governed by structured flower surface is vital for successful pollination^25^.

Floral reflectance spectrum was unique to all species studied. The spread of floral spectral reflection within each region of spectrum indicated by its skeweness points to the silent efforts made to attract pollinators. Differences in selective spectral absorption and consequent coloration were discussed by Sir C V Raman^6-9^. Nature has evolved flowers not to please man. It is a means to ensure plant evolution. Flower colors that we see around have evolved under selection by pollinators^2&5^. It will be erroneous to classify flower colors as perceived by human beings, because major pollinators have evolved themselves into a different visual system with sensitivity to UV, blue and green^4^.

## 4. Conclusion

This paper describes floral spectral reflectance of seven visibly distinguishable flowers of Trivandrum and introduces floral radiometry as a sub-discipline of ecology in India. Discrepancy between floral reflection and color perceived by humans is distinct from the results. Floral spectral reflectance is unique to a species. It provides a measurable biophysical parameter to characterize landscape level changes.

## 5. Acknowledgement

Authors thank Dr. Rama Rao N, Associate Professor, IIST Trivandrum for providing spectroradiometer used in field measurement. We also thank Director, IIITM-K and Government of Kerala for facilitating and supporting this study.

